# Attentional selection and communication through coherence: Scope and limitations

**DOI:** 10.1101/2023.08.16.553483

**Authors:** Priscilla E. Greenwood, Lawrence M. Ward

## Abstract

Synchronous neural oscillations are strongly associated with a variety of perceptual, cognitive, and behavioural processes. It has been proposed that the role of the synchronous oscillations in these processes is to facilitate information transmission between brain areas, the ‘communication through coherence,’ or CTC hypothesis.The details of how this mechanism would work, however, and its causal status, are still unclear. Here we investigate computationally a proposed mechanism for selective attention that directly implicates the CTC as causal. The mechanism involves alpha band (about 10 Hz) oscillations, originating in the pulvinar nucleus of the thalamus, being sent to communicating cortical areas, organizing gamma (about 40 Hz) oscillations there, and thus facilitating phase coherence and communication between them. This is proposed to happen contingent on control signals sent from higher-level cortical areas to the thalamic reticular nucleus, which controls the alpha oscillations sent to cortex by the pulvinar. We studied the scope of this mechanism in parameter space, and limitations implied by this scope, using a computational implementation of our conceptual model. Our results indicate that, although the CTC-based mechanism can account for some effects of top-down and bottom-up attentional selection, its limitations indicate that an alternative mechanism, in which oscillatory coherence is caused by communication between brain areas rather than being a causal factor for it, might operate in addition to, or even instead of, the CTC mechanism.

**Author summary:** The ability to select some stimulus or stimulus location from all of those available to our senses is critical to our ability to navigate our complex biological and social niche, and to organize appropriate actions to accomplish our goals, namely to survive and to reproduce. We study here a possible mechanism by which the brain could accomplish attentional selection. We show that the interaction between 10 Hz and 40 Hz neural oscillations can provide increases in coherence and information transmission between computationally modelled cortical areas. Our results support the idea that synchronization between oscillations facilitates communication between cortical areas. Limitations and sensitivities to the model, however, also point to the idea that the relation between coherence and communication could go in the opposite direction in some cases, particularly when attention is not involved.

## Introduction

Ever since the discovery of stimulus-specific, synchronized, neuronal oscillations, in both spiking and local field potentials (LFPs), in the gamma band (around 40 Hz) in the visual cortex of the cat [1], there has been speculation about the role of these oscillations in the brain. Of particular relevance to the present paper is the suggestion that information flow in the cortex could be facilitated by synchronization of the oscillations between different cortical areas [2–4]. This suggestion has been buttressed by the work of many researchers, e.g., [5–11] to cite only a few, who showed behaviourally-relevant synchronization in brain activity at various scales of time and space. The idea that synchronized neuronal oscillations could facilitate communication between various brain areas has come to be called ‘communication through coherence’ (hence CTC) [12, 13]. Many empirical studies have provided circumstantial evidence that CTC is indeed importantly involved in how information is transmitted between brain areas, e.g., [14–17] to cite only a few, and also c.f. [18]. Thus, CTC as a theoretical idea has seemed to be on a relatively firm footing empirically, although definitive causal evidence is still lacking.

Importantly, though, recently CTC has been challenged as getting the causal order wrong. Instead, some work suggests, the order should be ‘coherence through communication’ (CTC’). In other words, neuronal coherence is said to be an epiphenomenon of communication between brain areas, rather than a facilitator of that communication. In particular, Schneider and colleagues [19] have created a model, and run experiments, showing that simply having connections between two brain regions allows bursts of spikes in a sending area to create similar bursts in a receiving area, as well as coherence between the local field potentials (LFPs) in the beta band (around 20 Hz) in the two areas. Interestingly, in the experiments, the receiving area did not display significant oscillations in the beta band, even while displaying LFP coherence with the sending area in the beta band, nor did the receiving area display phase locking between spikes and oscillations. Notably, their Synaptic and Source-Mixing (SSM) model, which does a good job of explaining their empirical results, does not involve coupling of oscillators, nor does it involve any kind of phase-locking between the sender’s and receiver’s spiking activity. In the SSM model, coherence and Granger causality between the sending and receiving areas depends on connectivity (number of active synapses) between sender and receiver, power spectrum of the sender (where in the spectrum power is relatively high), and the coherence between the LFP at the sender and the signal sent by the sender. Schneider and colleagues [19], in reviewing other relevant evidence, also suggested ‘Likewise, changes in inter-areal coherence with cognition or behavior should be very carefully interpreted …’ (p. 14).

In this paper we investigate the CTC concept in the context of a theoretical and computational model of a mechanism by which selective attention could choose salient signals from among the many impinging on an organism. This model enables a convincing account of this mechanism, as well as some important limitations to which it is subject. These limitations point to the reverse model, CTC’, as possibly partially replacing, or operating along side, the original CTC model. One way in which both of the two mechanisms could be operating in the brain is as follows: the attentional selection mechanism operates using CTC to enhance information transmission for attended stimuli, whereas uninvolved brain areas typically operate as described by the CTC’ concept, according to the SSM model of [19] or something like it, when communicating information that is not attended and/or does not reach consciousness.

### Phase difference and attention

It is well known that voluntary orienting of attention to a specific location, or to a specific feature or object, enhances processing of that location or stimulus, e.g., [22]. One way in which this could work is for there to be enhanced information transmission about the attended stimulus between the cortical areas that are processing the sensory information. In the light of the CTC hypothesis, it is then reasonable to suggest that such enhanced information transmission could result from synchronization of neural oscillations between the relevant cortical areas. A study by Saalman and colleagues [23] (see also, more recently, [24]) reported that oscillations in the alpha band (around 10 Hz), originating in the pulvinar nucleus of the thalamus, were closely tied to top-down attentional processing in the cortex. Taking this suggestion seriously, Quax and colleagues [20] created a spiking neuron network model of two communicating cortical areas that received sine-wave alpha input as if from the pulvinar nucleus. They showed that the alpha input organized the gamma oscillations in the cortical areas, increasing both coherence and information transmission between the areas, consistent with the CTC hypothesis. Surprisingly, the alpha inputs to the two cortical areas in their study had maximum effects for the specific phase difference of Δ*ϕ* =*−π/*2 between the 10 Hz oscillations sent to the two cortical areas.

Intrigued by this finding, Greenwood and Ward [21] wondered what was special about *−π/*2. They proved a lemma that showed that, in fact, phase coherence between summed sine waves, say *α*_1_ + *γ*_1_ and *α*_2_ + *γ*_2_, similar to those in the study of [20], was maximum when the phase difference in radians, Δ*ϕ*, between *α*_1_ and *α*_2_ equalled that between *γ*_1_ and *γ*_2_, i.e., in the case of the results of [20], *−π/*2. The finding of [20], that maximal effects of alpha inputs to gamma-oscillating cortical areas were found when Δ*ϕ* = *−π/*2, pointed to the possibility that there was a similar phase difference, Δ*ψ*, between the gamma oscillations in the two modelled cortical areas. Greenwood and Ward then showed that the results of [20] could be replicated using a different macro-model of cortical neural activity to generate the gamma oscillations while retaining the noisy sine wave input for the alpha oscillations. Their rate model is similar to that originated by Wilson and Cowan [25] that, with noise, produces quasi-cycles. Quasi-cycles are noisy periodicities resulting from damped oscillations sustained by noise, e.g., [26]. In a separate part of their study, Greenwood and Ward implemented a similar rate model for the 10 Hz oscillations emanating from the pulvinar, replacing the noisy sine waves used by [20] with quasi-cycles. Their replication of the results of [20] using this method to produce oscillations completed the expression of the cortex-pulvinar model entirely in terms of quasi-cycles.

Fig 1 demonstrates the effects of alpha-gamma phase offset matching on phase coherence, maximum signal onset effect, and mean signal onset effect around the maximum, from the study of [21]. Clearly the maxima of these measures occur when the phase offset of the alpha waves approximately matches the phase offset of the gamma waves, at *−π/*2 in the top part of Fig 1, and near 0 in the bottom part of Fig 1. As mentioned, these results closely mirror those of [20], but with quasi-cycle oscillations rather than populations of spiking neuron models and sine waves. These results of [21] suggest that it is the oscillations that are essential in this situation, rather than the details of the neuron spiking that causes the oscillations. The rate model in [21] has no spiking, only oscillations, but retains the results of [20].

**Fig 1.**
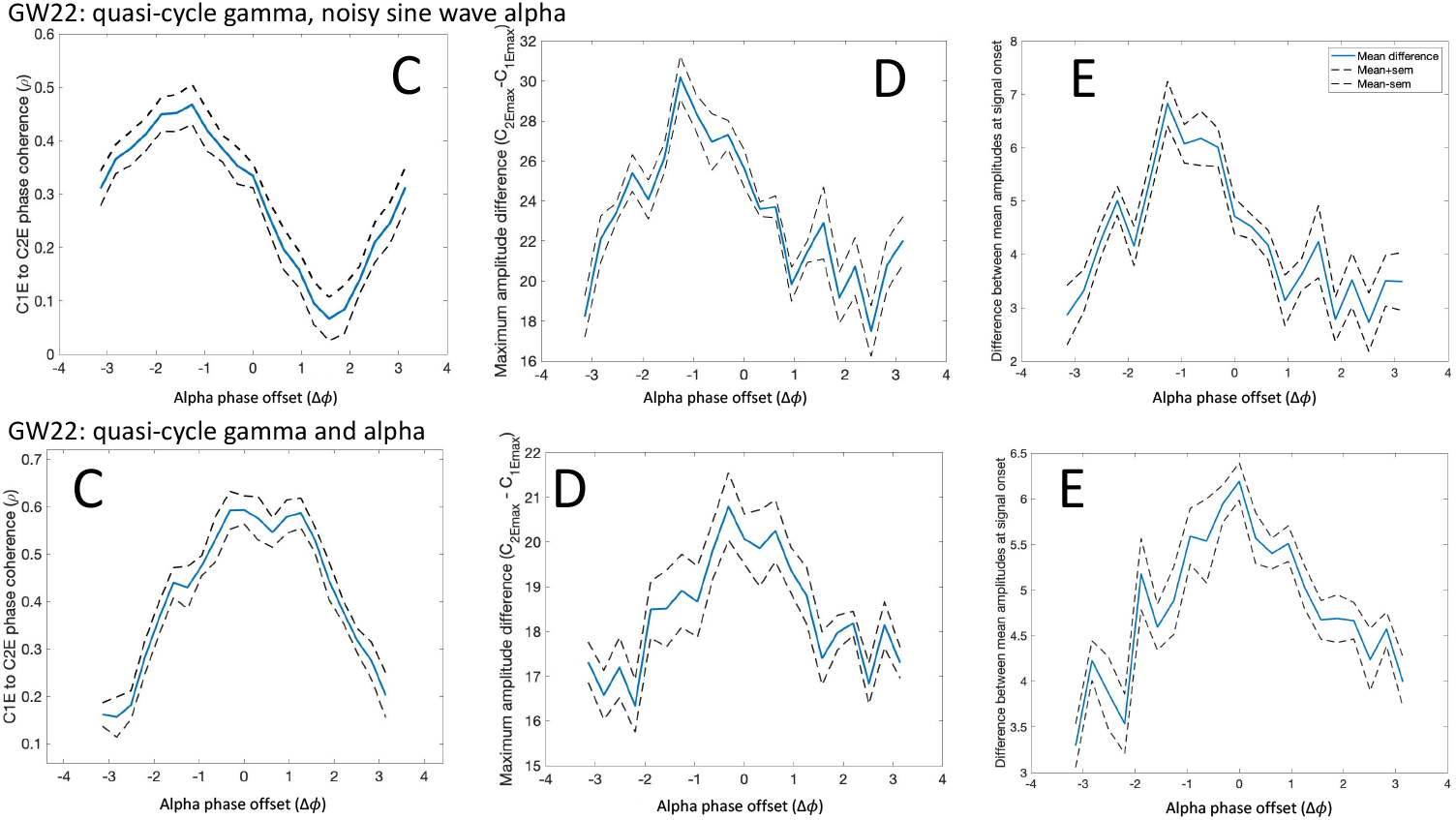
Results of [21] demonstrating the effects of phase offsets of alpha and gamma oscillations on (C) phase coherence, (D) maximum signal onset effect, and (E) mean signal onset effect around the maximum. The top part of the figure, ‘GW22 quasi-cycle gamma, noisy sine wave alpha,’ displays results from Fig 3 of [21] where the gamma oscillations were quasi-cycles and the alpha oscillations were noisy sine waves. The bottom part of the figure, ‘GW22 quasi-cycle gamma and alpha,’ displays results from Fig 4 of [21] where both gamma and alpha oscillations were quasi-cycles. The *gamma* phase offsets (not shown) are different for the two cases: *−π/*2 for the results in the top part and 0 for the results in the bottom part, respectively. Thus, the maximum phase coherence and signal onset effects occur near *−π/*2 in the top part and near 0 in the bottom part. Reproduced from [21] with permission.

Another recent study has also implicated phase differences between oscillations as important in information transfer between cortical areas [27]. In a context similar to that of [20], communicating populations of model neurons were simulated for two cortical areas. Here the frequencies of the oscillations in the sending and receiving areas, as well as the time delay (translated to a phase delay) between them, were varied.

Again, in this study, an optimal phase difference was discovered; in this case information transmission and coherence were maximum at a phase delay of *π* between the oscillations in the two areas. In the light of these additional findings, we undertook to elaborate the model studied by [20, 21], aiming to demonstrate both its generality and usefulness as well as to reveal its limitations.

### Selective attention via phase offset

Here we describe an elaborated model of attentional selection similar to that studied in [20, 21], specifically controlled by pulvinar input to communicating cortical areas (here assumed to be V1 and V2 in the visual system) as in [23, 24, 28]. The model operates to focus attention on one out of several stimuli that are present at once, whether in a ‘top-down,’ voluntary, fashion or in a ‘bottom-up,’ involuntary, fashion, as attention does in the brain [22]. Fig 2 displays a conceptual picture of this model, which is an elaboration of Fig 2C in [21]. Within Fig 2 are two copies of Fig 2C in [21], with parts labelled X and Y. The two parts appear as separate if the inputs, signals X and Y, and the TRN are removed, although in actual cortex there might be connections between them. The two copies are representatives of the many ”channels”, or subsystems, in each sensory cortical region that are tuned to different features of stimulus inputs, including in vision, e.g., position in space, color, shape, and so forth. So, in Fig 2, channels X and Y represent groups of neurons in V1 that respond to the features of signals X and Y, respectively. The elaboration consists of the addition of circuitry involving the Thalamic Reticular Nucleus (TRN) and its interaction with the pulvinar generators of 10 Hz oscillations.

**Fig 2.**
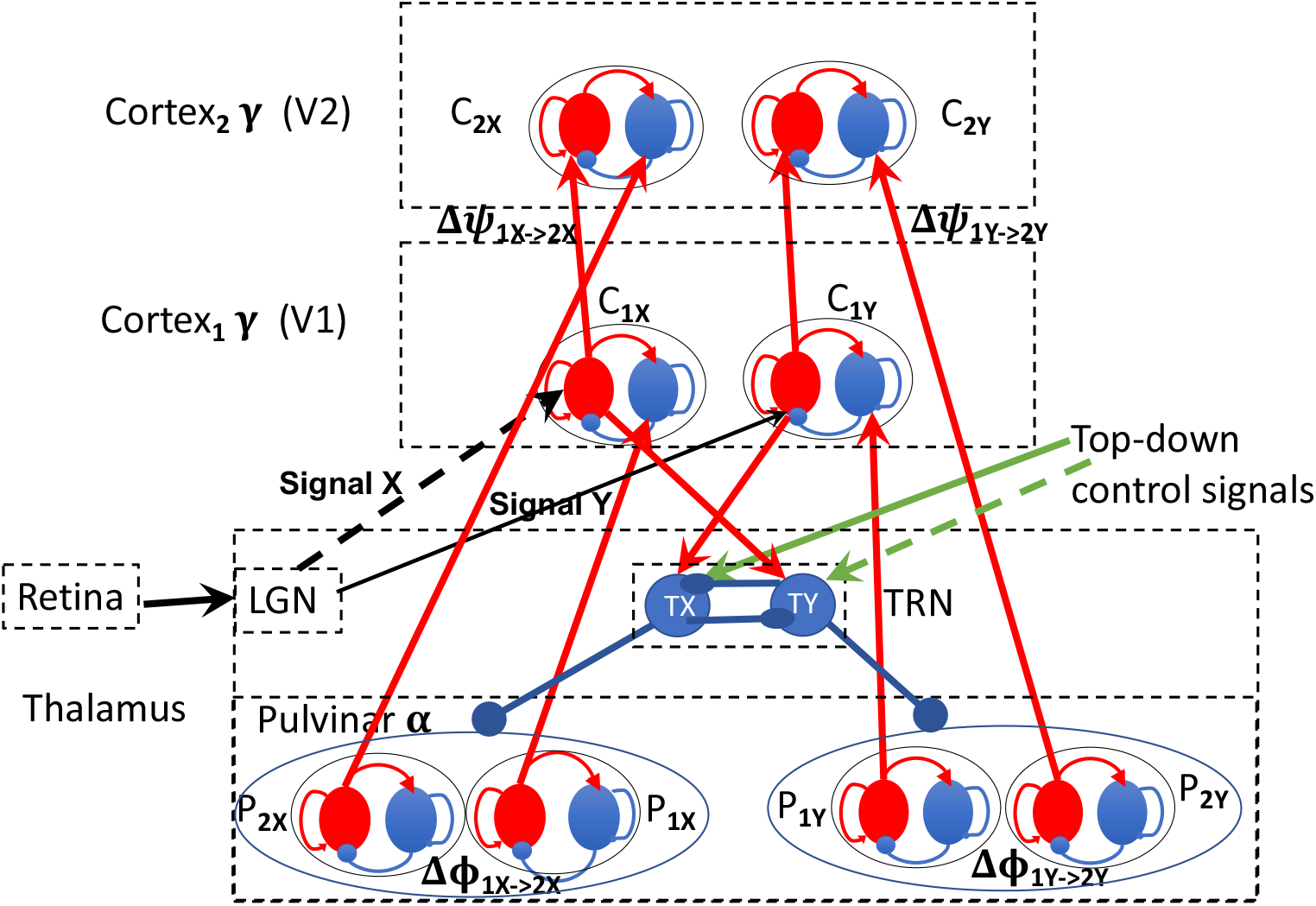
Structural model of top-down and bottom-up attentional selection. EI pairs are red (excitatory, E) and blue (inhibitory, I) disks; red arrows are excitatory, blue arrows are inhibitory, black arrows represent input signals, or stimuli, - they add increments in neural activity to the processes to which they connect. Top-down signals, green arrows, excite thalamic reticular nucleus (TRN) inhibitory neurons in the same way as do inputs from V1. The (visual) signals excite the V1 neurons that are tuned to them, e.g., vertical (say, signal X) or horizontal (say signal Y) lines on the retina excite V1 neurons tuned to vertical or horizontal lines respectively via tuned neurons in the LGN.

The elaborated system displayed in Fig 2 operates in top-down mode as described in [20, 21]. First, as in [20, 21], it is assumed, and implemented by parameter choices, that cortical areas oscillate at approximately 40 Hz, and pulvinar inputs to cortex oscillate at approximately 10 Hz. In the model of [21] both oscillations are produced internally to cortex and pulvinar as noisy quasi-cycles as defined in (1), although other generators of neural oscillations, such as PING models, are not being ruled out. Next, it is assumed that the phase offset, Δ*ψ* between V1 and V2 gamma oscillations, arising from the synaptic delay between V1 and V2 neurons, is fixed at the same value for all signals to which V1 neurons are tuned. The phase offset, Δ*ϕ*, of the 10 Hz pulvinar inputs to V1 and V2 is fixed at approximately the same value as the phase offset between V1 and V2. Thus, the phase offsets between alpha and gamma oscillations usually approximately match, meaning, as shown in [20, 21], that there is relatively high coherence and information transmission between V1 and V2 when corresponding areas of V1 and V2 receive 10 Hz oscillations from the pulvinar. The TRN contains only inhibitory neurons and is connected to all of the 10 Hz oscillators in the pulvinar. Thus, the TRN can modulate alpha input to V1 and V2 by inhibiting the output of particular 10 Hz oscillators to V1 and V2. For example, in Fig 2 each of the two signals, X and Y, will excite their respective E processes of *C*_1*X*_, *C*_1*Y*_ in V1, which in turn excite both their corresponding E processes *C*_2*X*_, *C*_2*Y*_ in V2, and their opposite inhibitory processes in TRN, TY and TX. The inhibitory processes in TRN inhibit each other as well [29], so there is a competition between incoming stimuli implemented in the TRN. Exciting the inhibitory TRN processes will inhibit activity in the relevant pulvinar neurons, *P*_1*Y*_ and *P*_2*Y*_ for signal X and *P*_1*X*_ and *P*_2*X*_ for signal Y, to the extent permitted by the internal TRN competition and any top-down control signals.

In a typical top-down visual attention scenario, there would be a variety of stimuli available in the visual scene, say X and Y. One of these, at some location and extent in the visual field, would be selected based on salience given by current goals, previous interactions, memory, etc. Top-down control signals implementing ‘voluntary’ attentional selection (re these goals, search targets, etc.) would affect the TRN neurons by strongly exciting those TRN neurons connected to cortical neurons tuned to stimuli *other than the desired stimulus*. In Fig 2 the desired stimulus initially is signal Y (via solid green arrow). Thus, the TRN would inhibit the 10 Hz pulvinar oscillations from going to those (unselected) tuned neurons in V1, *C*_1*X*_, that are responding to signal X. This would allow the 10 Hz and 40 Hz phase-offset match to induce optimal coherence and signal transmission between *C*_1*Y*_ and *C*_2*Y*_ in V1 and V2 for the selected signal Y. If signal Y is selected in this way, then phase coherence and signal transmission would remain suboptimal for signal X and any other signals because of lack of 10 Hz input, and signal Y would be ‘attended.’

Competition between TRN neurons would mediate between signals of different strengths and top-down selected signals for precedence, as well as between signals in different sensory modalities (and primary sensory cortical areas). Top-down attention can overcome some differences in signal strength as depicted in our example (signal X arrow is thicker, e.g., [31]). Some signals, however, are simply too strong, e.g., a very bright flash of light, say signal X suddenly flashes brightly, indicated by the X line being dashed. In this case the the bottom-up, ‘involuntary,’ or ‘automatic,’ mechanism might pull attention away from whatever top-down signal was being selected at the time [22]. We assume that, as in behavioural studies, this pulling of attention to a new stimulus is followed immediately by reorienting top-down attention to the new stimulus until its relevance is ascertained. Even though attention currently is being paid to signal Y through the top-down mechanism just described, the brightening of signal X will have several effects. First, it will produce a very strong signal onset effect, in spite of the fact that no pulvinar 10 Hz oscillation is being sent there because of the allocation of top-down attention to signal Y (see *Results* for a supporting simulation result). This will be transmitted to executive areas of the brain responsible for sending signals to implement top-down attention. Also, because the very strong signal X wins the competition between the TRN neurons, and this will be reinforced by a top-down signal exciting TY (indicated by the dashed green arrow), 10 Hz input from *P*_1*Y*_ and *P*_2*Y*_ to *C*_1_*Y* and *C*_2_*Y* will be terminated. Thus, 10 Hz oscillations from *P*_1*X*_ and *P*_2*X*_ now will be injected into the 40 Hz oscillating *C*_1_*X* and *C*_2_*X* in V1 and V2, respectively. The roughly equal phase offsets between 10 Hz and 40 Hz oscillations for signal X now will optimize the phase coherence and signal transmission between V1 and V2 for signal X, as described in [21]. Because the 10 Hz input from *P*_1*Y*_ and *P*_2*Y*_ to *C*_1_*Y* and *C*_2_*Y* has been blocked by TY, however, the coherence and signal transmission between relevant cortical areas become lower (but not 0) for signal Y than for signal X. The large signal onset effect of signal X at V2 (and likely further up the visual system), is presumed to be what it means for signal X, the very strong signal, to ‘pull,’ or ‘draw,’ attention to itself in a bottom-up manner. There is a wealth of evidence that this effect is ubiquitous, both in the brain and in behaviour [22].

Our contribution here is the above description of a possible mechanism for the interaction between the pulvinar, the TRN, and the cortex, to select an object of attention. The question arises: to what extent is the mechanism dependant upon oscillations? Our conception of an implementation is in terms of quasi-cycles, which we now introduce.

## Methods

### Computational implementation of the model

As in [21], here we use an EI rate model that produces quasi-cycle oscillators that are described in detail in [30] based on a model of [32]:

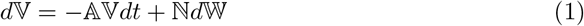

where

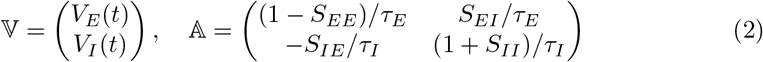

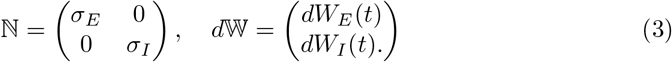

This model is a description of the time evolution of the local field potentials (LFPs), closely related to firing rate, of two coupled populations of neurons, excitatory, *V*_*E*_(*t*), and inhibitory, *V*_*I*_ (*t*). The *V*_*E*_ and *V*_*I*_ processes defined by (1), (2), (3) will be called the E and I processes. We refer to a stochastic process defined by a model of the form (1) as an EI pair. The parameters in 𝔸 represent the synaptic coupling strengths, *S*_*ij*_, and time constants, *τ*_*i*_, between the excitatory and inhibitory neuron populations, *σ*_*i*_ are noise amplitudes, and *W*_*i*_ are standard Wiener noise processes. In this model, when the eigenvalues of *−*𝔸 are complex, *λ i ω*, with 0 *< λ << ω*, the system 1 has sustained (noisy) oscillations, called quasi-cycles, in a narrow band around *ω*. The parameters in *−*𝔸 can be adjusted to give a wide range of quasi-oscillation frequencies within the range relevant to brain activity related to mental processes [30]. The parameter values used in 𝔸 to produce approximately 40 Hz oscillations in the EI pairs are listed in Table 1. The left hand side of the diagram in Fig 2, without the TRN or signals (or, similarly, the right hand side by itself), was the subject of [21]. Simulations in [21] used (1) to produce independent quasicycles of central frequency gamma in the EI pairs denoted *C*_1*X*_, *C*_2*X*_, and of central frequency alpha in the EI pairs denoted *P*_1*X*_, *P*_2*X*_ in Fig 2.

**Table 1.**
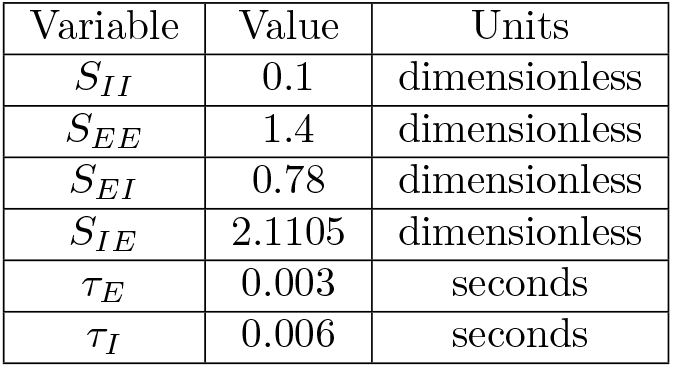
Parameters of 𝔸 used in 40 Hz EI pair simulations.

| Variable | Value | Units |
| --- | --- | --- |
| $S_{II}$ | 0.1 | dimensionless |
| $S_{EE}$ | 1.4 | dimensionless |
| $S_{EI}$ | 0.78 | dimensionless |
| $S_{IE}$ | 2.1105 | dimensionless |
| $\tau_E$ | 0.003 | seconds |
| $\tau_I$ | 0.006 | seconds |

In this paper we have opted to implement the elaborated model in the simplest possible way, given that the main effects have already been demonstrated in both populations of individual spiking neuron models [20] and of generic rate models like (1) [21]. Both types of models emphasize periodic alpha and gamma oscillations, although the population models exhibit the oscillations in bursts of spiking organized by a sinusoidal input, whereas the rate model exhibits interacting quasi-cycles. The process defined by (1) can be factored into a rotation (sinusoid) multiplied by a 2D Ornstein-Uhlenbeck process [26], and so is essentially a noisy sinusoid.

Each of the two signal channels in the model of Fig 2 is computed as follows. The channels of the cortical processes (V1 and V2) are modelled as

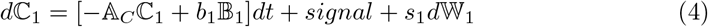

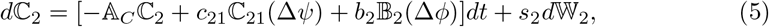

where 𝔸_*C*_ is the 𝔸 matrix of coefficients that generates 40 Hz oscillations, damped but sustained by noise, ℂ_1_, ℂ_2_ are 2D vector processes, (*C*_1*E*_, *C*_1*I*_)′, (*C*_2*E*_, *C*_2*I*_)′, for EI pairs, as in (1), (2), (3), oscillating at natural frequency 40 Hz but with added 𝔹_1_ = (0, *−α*_1_(*t*))′, 𝔹_2_(Δ*ϕ*) = (0, − *α*_2_(Δ*ϕ, t*))′, *b*_1_, *b*_2_ are the amount of alpha injected into *C*_1*I*_, *C*_2*I*_. Further, ℂ_21_(Δ*ψ*) = (*C*_1*E*_(*t* + Δ*ψ*, 0)′ represents the input to *C*_2*E*_ from *C*_1*E*_ delayed by Δ*ψ*, where Δ*ψ* is the phase difference between *C*_1*E*_, *C*_2*E*_ caused by adding a fraction, *c*_21_, of *C*_1*E*_ to *C*_2*E*_ at a time delay that implements the phase offset between V1 and V2. The *signal* in (4) is modelled as a constant input, i.e., a step function, as in [20, 21], except where indicated. Although common in both in vivo and in vitro studies, this is problematic because LGN input to V1 actually consists of structured bursts of spikes [33]. This limitation will be discussed in *Sensitivities and Limitations*.

As we demonstrated in [21], modelling the pulvinar 10 Hz inputs as noisy sine waves or as EI pair quasi-cycles gives rise to similar results, so here we use noisy sine waves for simplicity and to enable the most precise phase offset of the alpha waves. The inputs from the noisy sine waves oscillating at approximately 10 Hz are:

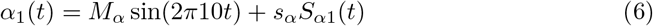

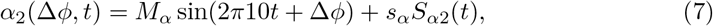

where *M*_*α*_ is the amplitude of the alpha sine waves and Δ*ϕ* is the *α*_1_ to *α*_2_ phase difference. The white noise, *S*_*α*_(*t*), added to the alpha sine waves has standard deviation *s*_*α*_.

Sample paths of processes ℂ_1_, ℂ_2_, defined by equations (4), (5) were simulated using the Euler-Maruyama algorithm, with the current values of the noisy 10 Hz sine waves added to the SDE at each time point.

Table 2, which is the main focus of this paper, displays the results of a large number of simulations with strategically chosen representative values for the parameters of Equations (4), (5), (6), (7). The simulations were run for 100,000 time points at 0.00005 sec per time point, so for 5 sec. A signal was added at time point 50,000 and remained on until time point 100,000. As in [21], phase coherence, *ρ*, was calculated as 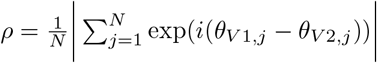, where *θ* is the instantaneous phase of the V1 or V2 oscillation computed from the analytic signal via the Hilbert transform, and *N* =number of time points over which *ρ* is computed. Here *ρ* was computed over *N* =100,000 time points. Signal onset effects, as in [20, 21], were measured as the maximum (max) of the positive part of the oscillation in V2 minus that in V1 between time points 50,000 and 55,000 (250 ms after signal onset), and the mean positive part of the oscillation over the interval max-1000 and max+1000 timepoints (50 ms).

**Table 2.**
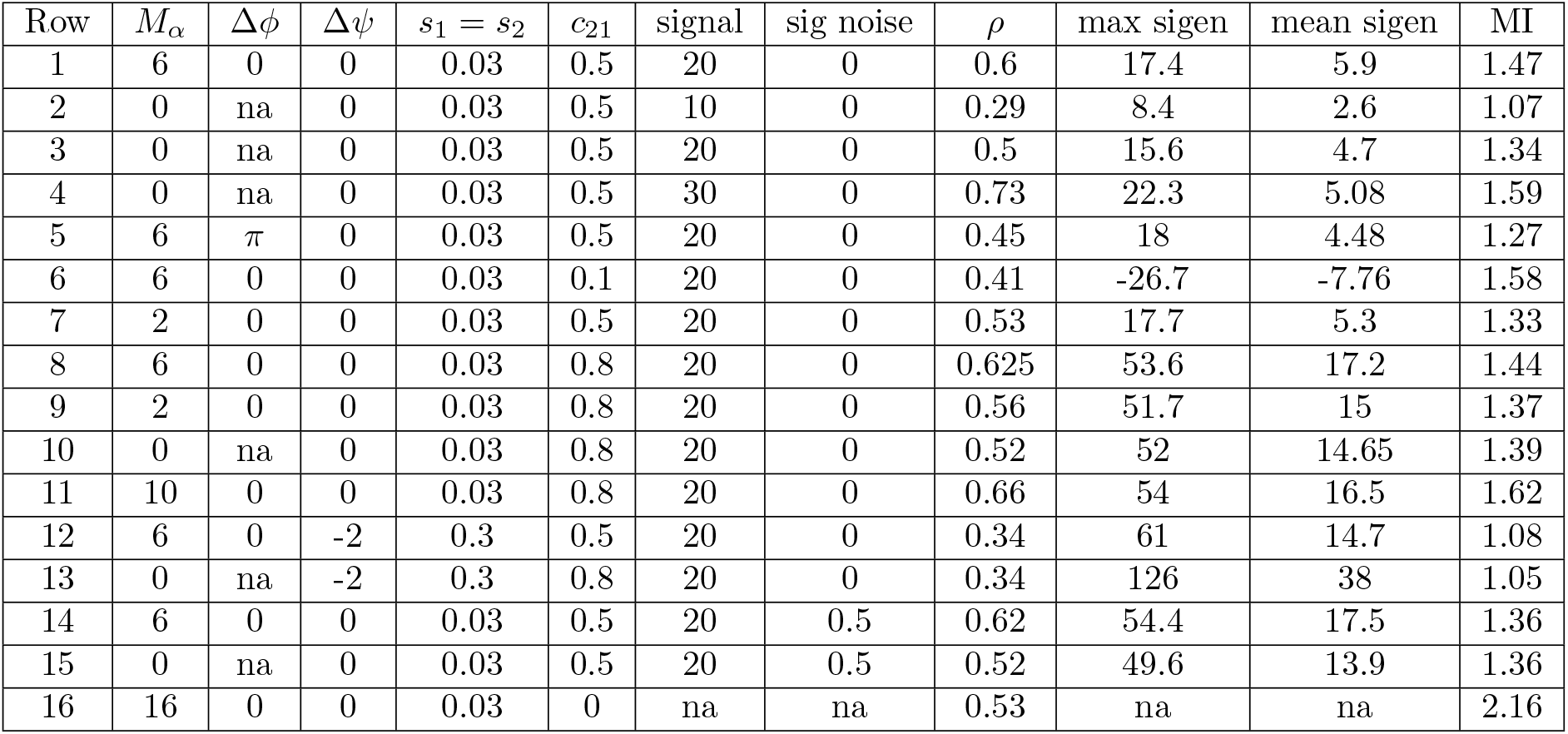
Results of simulations of model in Fig 1 using parameters of (4),(5),(6),(7). Parameter values are in the leftmost seven columns, output values in the rightmost four columns. “sig noise” is signal noise; “max sigen” is maximum of the positive part of the oscillation at V2 minus that at V1, i.e., *max*(*V*_2+_ *− V*_1+_), between signal onset and signal onset + 250 ms; “mean sigen” is the mean of the positive part of the oscillation between time at maximum - 50 ms and time at maximum + 50 ms at V2 minus that at V1; MI is mutual information between the phases of V1 and V2 after signal onset. All output values, *ρ*, max sigen, mean sigen, and MI, are the means of 10 realizations, with standard deviations approximately 0.01 to 0.02.

Mutual information, *MI*, was computed between V1 and V2 using a Matlab function [34] as

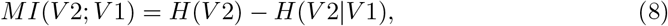

where *H*(*V* 2) is the entropy of V2, and *H*(*V* 2 │*V* 1) is the entropy of V2 conditional on V1. MI was computed separately for the interval before signal onset and for the interval after signal onset, for the raw time series of V1 and V2, and also for their phases and amplitudes. Phases and amplitudes were computed from the analytic signal and the Hilbert transformed time series. As the results for the various MI computations were similar, only those for phase-based MI for the interval after the signal onset are reported here.

## Results and Discussion

### Attentional selection mechanism

The first seven rows of Table 2 replicate the results of [21], and illustrate how the model of Fig 2 works for the bottom-up and top-down attentional selection scenarios. Row 1 is an example where a strong signal, *X* = 20, wins the competition in the TRN and thus prevents *α* oscillations from the pulvinar from reaching V1 and V2 neurons tuned to the weaker signal, *Y* = 10. For the stronger signal, *α* oscillations at the matching phase offset, 0 in this case, would reach the areas of V1 and V2 that respond to that signal, causing a fairly large phase coherence value *ρ* = 0.6, a strong signal onset enhancement of 17.4, and also a strong mean signal enhancement effect of 5.9. These values are consistent with those for the matching phase offset case in [21] (see Fig 1). For the weaker signal, Y, however, no *α* oscillations from the pulvinar reach V1 or V2 because of inhibition from the TRN. This situation is portrayed in Row 2 of Table 2. There, all but the signal strength being the same, V1-V2 phase coherence, *ρ* is substantially lower, as are the maximum and mean signal enhancement effects, consistent with both [20] and with [21]. Mutual information between V1 and V2 during the signal period is also lower when there are no *α* oscillations sent to V1 and V2 for signal Y. Making Δ*ϕ* = *π* ≠ Δ*ψ* = 0, on the other hand, even with the fairly strong signal X=20, reduces both *ρ* and MI and the mean signal enhancement effect (Row 5), similar to results of [20, 21] (see Fig 1). Note that reducing the *magnitude, M*_*α*_, of the *α* oscillations to 2 from 6, as in Row 7, has only a small effect on *ρ* and mutual information, and no effect on signal enhancement relative to Row 1 values. So at least some input from the pulvinar would seem to be sufficient for the attentional effects we see.

Row 3 compared with Row 1 of Table 2 shows the effects of top-down attention selecting one of two signals. If the two signals have equal magnitude, then the selected one (Row 1) still shows greater phase coherence, signal onset effects, and mutual information than does the unselected one (Row 3), again because the TRN has been signalled to inhibit *α* coming from the area of the pulvinar connecting to the unselected neurons in V1 and V2.

On the other hand, if a very strong signal, say X=30, suddenly appears, attention can be pulled away from its current focus by very large signal onset effects, as in Row 4, despite the fact that, at the moment the strong signal appears, no pulvinar 10 Hz oscillations are going to the neurons tuned to the strong signal. Subsequently, however, 10 Hz oscillations would be sent to those neurons because of the mutual effects of the strong signal and top-down signal exciting TY, thus inhibiting *P*_1*Y*_, *P*_2*Y*_, and releasing *P*_1*X*_, *P*_2*X*_ from inhibition by TX.

Thus, our elaborated model uses the same basic mechanism, TRN inhibition of pulvinar *α* oscillations, to accomplish both top-down and bottom up attentional selection. In the next section we will see that the results displayed in Table 2, Rows 6-16, point to sensitivities and limitations of our model.

### Sensitivities and Limitations

In evaluating a model such as the one studied here, one is interested in its behaviour not only at the optimal parameter values, but also elsewhere in the parameter space. A useful model will be sensitive to parameter changes in a way that mirrors the behaviour of the phenomenon itself, while not oversensitive to small changes to which the phenomenon would be robust. In our present case these questions present difficult challenges, but we can make several comments based on the findings in Table 2. Not surprisingly, however, the ”success” of a model depends on the balance of several effects. When balance fails, pathologies are known to arise. Some of the sensitivities we are about to discuss could point to a source of pathologies in attention, or in neural transmission more generally.

Our elaborated model can account for basic attentional phenomena, in particular the association of phase coherence with information transmission between neural areas, as in the CTC hypothesis. There are, however, important limitations to the model, which we will illustrate using Table 2. Some are obvious: we are modelling at a macro scale, our EI equations are rate approximations, parameter values are not derived from empirical data, the details of both cortical and thalamic (TRN and pulvinar) connectivity are missing, and so forth. Importantly, it seems we and others have found a sweet spot in parameter space within which the indicated mechanism operates. But parameter values outside that sweet spot yield a somewhat different picture. For example, the connectivity between V1 and V2 is critical. Reducing the connectivity from 0.5 to 0.1, as in Row 6 of Table 2, yields moderately high phase coherence (0.41), but *negative* signal onset enhancement and *increased* mutual information. Increasing the V1-V2 connectivity to 0.8 while leaving everything else unchanged, as in Row 8, leads to values of *ρ* and MI comparable to those for Row 1, but accompanied by much larger values for signal enhancement. Finally, increasing the amplitude of the *γ* quasi-cycles by a factor of 10, to 0.3, as in Row 12, leads to a relatively small value for *ρ* and MI, but a large value for signal enhancement. Abolishing the *α* input under those conditions, Row 13, gives rise to a huge value of signal enhancement but still accompanied by relatively low *ρ* and MI. Interestingly, in both of these latter cases, the centroid of the distribution of phase differences between V1 and V2 was shifted to -2 radians, not matched to the 0 phase offset of the *α* frequency. These values are clearly inconsistent with CTC. They are also inconsistent with the elaborated model we have described here, in which offset-matching of 10 Hz input to cortex is the most important factor in affecting signal transmission between cortical areas.

Another limitation of both our and other models is the question of how input signals (stimuli) should be modelled. Some authors, e.g., [20, 21], have used a constant, i.e., step function, signal. Others have used a burst of Poisson or Gaussian pulses (basically, noise), and still others have used a constant plus some noise. Some, e.g., [35], have used a signal modelled on actual output from the LGN as input to V1. It does appear, however, that using as input the output from a noisy integrate and fire neuron (with a constant input to the model neuron) provides a good approximation of LGN output statistics, and has been shown to give better orientation selectivity than does LGN output modelled as an inhomogeneous Poisson point process [33]. Use of a constant as our input signal represents another limitation of our and others’ model results. Interestingly, when we input a noisy constant to our EI modelled V1, as in Row 14 of Table 2, we see little difference in *ρ* or MI from Row 1, but a very large difference in the signal enhancement effect. Interestingly, the added noise both increases the amplitudes of the quasicycles in (4) and (5) and makes them noisier, which might be partially responsible for the mentioned effect. And even without *α* input, Row 15, these effects are maintained. So noise is apparently playing a very large role in the signal enhancement effect. Note that both our quasi-cycle model, here and in [21], and the neuron population model of [20], involved noisy (stochastic) processes.

#### Coherence with no connection

The CTC hypothesis assumes that the coherence between two cortical areas implies a direct connection between them, because information is being transmitted from one to the other. In other words, the spikes from the sending area must arrive at the receiving area at an optimal point in the phase resetting curve. But such a connection is not necessary for there to be phase coherence between two cortical areas. For example, Row 16 of Table 2 shows that when the V1-V2 connection is zero, but the *α* driving is large, high phase coherence is observed between V1 and V2, along with even higher MI. Of course, with no connection there is no information transmission between the areas. Such a result can mislead analysis of neural data for functional connectivity when several sources are simultaneously active but only simple pairwise phase coherence is measured. In these cases significant coherence between two sources could arise because each is coherent with a third source, or even multiple other sources. Most authors are aware of this problem, but conditional functional analysis, as in multiple linear regression, especially nonlinear analysis as in transfer entropy, is more complicated, and so is seldom done.

#### Coherence through communication

It is clear from the results displayed in Rows 8-11 in Table 2 that driving of one cortical area by another can result in high phase coherence and MI, as well as large signal onset enhancement effects, regardless of the magnitude of *α* input to them, including none at all (Row 10). This is the scenario envisioned by the CTC’ hypothesis espoused by [19] and also studied by [27]. As mentioned in the introduction, it is possible that this mechanism could be responsible for coherence between cortical areas when information is being transmitted outside of attention or consciousness, even while the pulvinar-origin *α* mechanism could be responsible for attentional enhancement of sensory inputs. Alternatively, it is possible that pulvinar-origin *α* input to cortex has some other function, somehow associated with attention but not actually causing phase coherence and enhanced information transmission.

#### Onset vs total signal effects

In the models studied by [20, 21] and here, the largest effects of *α* input to modelled cortical areas were at signal onset. As shown in previous sections, this is affected by phase offset matching but also by connectivity between areas (at least the way we compute it, as additive input) and by noise. There seems to be less effect on ongoing information transmission, at least as computed by MI over the entire time the signal is present. (We note that it is necessary to have a relatively long time series to compute MI in an unbiased way, as is also the case for phase coherence.) The smaller effect on ongoing signal transmission is advantageous in the context of a complex environment, allowing for other signals to interrupt with their own large onset effects or change in coherence/signal transmission.

#### Paradox re alpha

As we and Quax and colleagues have shown, alpha power *increases* in cortical areas that receive 10 Hz input from the pulvinar [20, 21]. However, many studies have shown that alpha power in EEG and MEG recordings actually *decreases* in the area of cortex where the attended stimulus is processed, and *increases* in the area of cortex where the unattended stimulus is processed, e.g., [11, 36, 37], although the details are not clear [38]. Moreover, alpha and gamma power seem to be anti-correlated, e.g., [10]. How can this be if alpha from the pulvinar is inhibited by the TRN for unattended signals, and allowed to go to V1 and V2 for attended signals, as in our suggested model? And what about the timing of the alpha power changes? In [11] alpha power (from MEG recording) first increased after a top-down cue, then decreased sharply around 400 ms after the cue in both cued and uncued hemifields, and then increased more in the ipsilateral (unattended) field at 500-700 ms post-cue (but did increase in both hemifields in that time range). This last time interval is also when narrow-band transfer entropy, similar to Granger causality, was significant between frontal and parietal areas. More recently, Hanna et al. [37] found similar results to [11]. They used phase transfer entropy with MEG recordings to show that alpha connectivity to, and alpha power in, unattended areas of sensory cortex (auditory or visual) decreased in a coordinated way. What is going on? This is a paradox.

One way to solve the paradox is to invoke locally-generated (i.e., in V1 and V2), but top-down signal stimulated or signaled, alpha for the power changes in unattended and attended cortex, e.g., [39, 40], recorded by EEG and MEG. Then pulvinar-generated alpha can be the source of the alpha that organizes the phase-offset-matched gamma oscillations that leads to good signal transmission [20, 21, 23, 24]. We are proposing that phase-offset-matching alpha input would be limited to attended cortex, with alpha input from pulvinar to unattended cortex suppressed via the TRN. Locally-generated alpha, however, could be increased in an unattended sensory area by changes in synaptic efficacies from values generating gamma to values generating alpha, resulting in less gamma, because the same EI pairs can’t generate both frequencies. This could happen on a millisecond timescale [41, 42]. This would mean it would be more difficult to process stimuli in an unattended area, because active processing is associated with increases in gamma power. Then 10 Hz from pulvinar could organize the gamma in the attended area, which is still generating gamma. Alpha originating in the pulvinar would not change (decrease) gamma-generating or gamma power in the attended area because synaptic efficacies wouldn’t be changed by input of this alpha. Alpha power would be increased somewhat by the alpha input from the pulvinar (but not to the level of the unattended cortex), however, and the gamma oscillations would be organized by it, as in [20, 21] and the present study, increasing signal transmission.

Another way in which the change in locally-generated alpha and gamma power could be implemented was described by Chen and colleagues [43], cf. also [44], and modelled by Domhof and Tiesinga [45]. Chen and colleagues found that changing the activities of two different types of inhibitory, GABAergic, interneurons, using optogenetic methods, dramatically affected the power in lower (5-30 Hz) and higher (20-80 Hz) frequency bands. In particular, inhibiting parvalbumin-expressing (PV) interneurons, which inhibit pyramidal neurons, increased gamma-band oscillations, whereas exciting somatostatin-expressing (SOM) neurons, which inhibit both PV and pyramidal neurons, increased lower-frequency oscillations. So, a top-down signal could accomplish the increase in local alpha power, while at the same time (same signal) decreasing gamma power, simply by exciting SOM neurons.

Whichever of these two mechanisms, or an unknown third, is responsible for the observed changes in alpha and gamma in unattended and attended areas of cortex remains to be determined. But it seems clear that some mechanism like one of the ones described must be operative to resolve the paradox in alpha oscillations during attention. Note that alpha oscillations are also suggested to play a role in timing of cognitive operations [46] and in long-range coordination in the brain [10], among other things. Thus, not only are alpha oscillations ubiquitous in the brain, but they may play a variety of roles in perception and cognition.

## Conclusions

We have elaborated a model of attentional selection in which alpha oscillations generated in the pulvinar nucleus of the thalamus organize gamma oscillations in cortical areas so that they are more coherent than without the alpha input. Thus, they transfer information about signals more efficiently than without that input. This model describes how attention can select a particular stimulus to emphasize, either though a top-down signal from other cortical areas, or by preferring more salient stimuli in a bottom-up fashion. There are significant limitations to the model as it exists, however. In particular, the model works well only in a limited parameter range, so it would be very sensitive to physiological conditions. Moreover, there is no consensus on how to characterize stimuli (or signals) in such models. Some researchers simply add a constant to the input, as is often done in physiological studies in vitro. Others, including us, have argued that signals should be inputs that increase the amplitude of the LFP, for example outputs of leaky integrate and fire neurons [33].

The relation between coherence and communication is not as simple as it has seemed from the CTC hypothesis. First, through input from a third area (e.g., the pulvinar), coherence can be increased even though there is no communication at all between the monitored areas. This can fool data analysis that only focuses on, e.g., phase coherence between pairs of cortical areas. Second, coherence between cortical areas can be increased simply by one area sending spikes to another: coherence through communication (CTC’). Third, it is not clear how to measure communication in these models. Some measure information transmission by focusing on the onset of the stimuli, whereas others focus on the entire time the stimulus is present, with somewhat different results.

Finally, it is becoming clear that alpha oscillations could perform many roles in the brain, from attention selection to timing to long-range coordination. It is important that these roles and their individual circumstances be clarified. Future modelling and empirical work might be able to determine when and how the different suggested mechanisms perform. But it will be important to remember the scope and limitations of these mechanisms. And also it is important to determine whether the ubiquitous neural oscillations play a causal role in any of these mechanisms, or simply arise epiphenomonally from the spiking of neurons as they transmit information throughout the brain.

## Acknowledgments

This research was supported by a Discovery Grant from the Natural Sciences and Engineering Research Council of Canada (NSERC) to LMW.

